# Intrinsic adaptive value and early fate of gene duplication revealed by a bottom-up approach

**DOI:** 10.1101/151910

**Authors:** Guillermo Rodrigo, Mario A. Fares

## Abstract

Gene duplication is a major source of functional innovations and genome complexity, albeit this evolutionary process requires the preservation of duplicates in the genomes for long time. However, the population genetic mechanisms governing this preservation, especially in the critical very initial phase, have remained largely unknown. Here, we demonstrate that gene duplication confers *per se* a weak selective advantage in scenarios of fitness trade-offs. Through a precise quantitative description of a model system, we show that a second gene copy enhances the information transfer from the environmental signal to the phenotypic response by reducing gene expression inaccuracies derived from pervasive molecular noise and suboptimal gene regulation. We then reveal that such a phenotypic accuracy yields a selective advantage in the order of 0.1% on average, which would allow the positive selection of gene duplication in populations with moderate or large sizes. This advantage is greater at higher noise levels and intermediate concentrations of the environmental molecule, when fitness trade-offs become more evident. Moreover, we show that the genome rearrangement rates greatly condition the eventual fixation of duplicated genes, either by natural selection or by random genetic drift. Overall, our theoretical results highlight an original adaptive value for cells carrying new-born duplicates, broadly analyze the selective conditions that determine their early fates in different organisms, and reconcile population genetics with evolution by gene duplication.

**SIGNIFICANCE:** Gene duplication is considered a major driver for the evolution of biological complexity. However, it is still enigmatic to what extent natural selection and genetic drift have governed this evolutionary process. This work uncovers a selective advantage for genotypes carrying duplicates, called phenotypic accuracy, widely characterized thanks to a multi-scale mathematical model coupling gene regulation with population genetics. Importantly, the integrative results presented here provide a detailed mechanistic description for the fixation of duplicates, which allows making predictions about the genome architectures, and which is relevant to understand the origins of complexity.

## INTRODUCTION

Gene duplication has enthralled researchers for decades due to its link to the emergence of major evolutionary innovations in organisms of ranging complexity (Ohno, 1970). The key aspect to deeply understand this process concerns the early stage, when the fate of the new-born gene is decided (Innan & Kondrashov, 2010). A classical theory predicts the fixation of duplicated genes in the population under neutral selective conditions (*i.e*., by random genetic drift; Kimura, 1983; Lynch & Conery, 2003). Hence, the loss of the new-born gene is the most common evolutionary fate. Once a duplicate is fixed, it is generally accepted that genetic redundancy leads to relaxed selection constraints over one or both gene copies, which increases the load in mutations (Lynch & Conery, 2000; Keane *et al*., 2014). In rare occasions, this evolutionary process leads to the origin of a novel, previously unexplored function by one of the gene copies (Conant & Wolfe, 2008).

However, because gene duplication imposes a cost to the cell by requiring additional resources for expression (Wagner, 2005; Lynch & Marinov, 2015; Price & Arkin, 2016), especially in simple organisms, and because it unbalances tightly regulated pathways that are instrumental for the cell (Papp *et al*., 2003; Birchler *et al.*, 2005), leading to diseases in complex organisms (Tang & Amon, 2013), purifying selection could preclude that fixation. A possible rationale that has been long recognized is that those duplicated genes that were fixed in the population immediately contributed with an adaptive value to the organism (Innan & Kondrashov, 2010). Even though, it is still stunningly unclear to what extent natural selection could also take part in the process that drives the fixation, and also initial maintenance, of duplicated genes according to population genetics (Lynch, 2007).

Two basic hypotheses have been proposed to explain the selective advantage of duplicated genes. First, a higher gene expression level resulting from duplication could be favorable (Riehle *et al*., 2001). This hypothesis requires that the ancestral system (pre-duplication) is far from the optimal operation point; as far as to assert that nearby 100% expression increase is beneficial. This seems plausible in extreme circumstances, but not in routine environments for which the organism should be adapted (King & Masel, 2007). It is then not surprising that many of the reported examples in which a greater gene copy number is favorable relate to sporadic, mainly stressing environments (Riehle *et al*., 2001; Gonzalez *et al*., 2005). Arguably, if a duplicate were fixed in one of these environments, it would be rapidly removed by purifying selection once the extreme circumstance ceased. Moreover, beneficial single-point mutations occurring in the *cis*-regulatory region of the gene of interest would be mostly sufficient to face several environmental changes (Wray, 2007). Thus, this model is insufficient to clarify the origin of most duplications, although it could explain some particular cases.

Second, the functional backup provided by the second gene copy upon duplication may allow the rapid accumulation of beneficial mutations, either to develop a novel function (Zhang *et al*., 1998; Bergthorsson *et al*., 2007), or to escape from the conflict of optimizing alternative functions (Hittinger & Carroll, 2007; Des Marais & Rausher, 2008). The positive selection of these mutations may of course occur, as suggested by the dN/dS values (> 1) reported for different genomic sequences (Han *et al*., 2009; Fischer *et al*., 2014). This requires, nevertheless, that the frequency of cells carrying a second gene copy in the population increases to a point at which a mutation in the duplicate is likely to be found; a condition that is not met during the critical very initial phase following duplication (Lynch *et al*., 2001). Therefore, such adaptive processes, although important for the long-term maintenance of duplicates, do not contribute much to increase their fixation probabilities.

In addition to these two hypotheses, it has been proposed that gene duplication could allow compensating for errors in the phenotypic response due to a loss of expression caused by genotypic or phenotypic mutations (Clark, 1994; Nowak *et al*., 1997; Wagner, 1999). This model needs to invoke high error rates to have an impact at the population level from the beginning, and then to reach prevalence of genotypes with duplication by overcoming genetic drift. Errors in phenotype could also be caused by stochastic fluctuations in gene expression (Elowitz *et al*., 2002), with gene duplication eventually reducing the amplitude of such fluctuations (Kafri *et al*., 2006; Lehner, 2010; Rodrigo & Poyatos, 2016). Despite, how (and how much) this strategy is really advantageous to invade a natural population, which includes of genetically and non-genetically heterogeneous individuals, and which continuously evolves under fitness trade-offs (Balázsi *et al*., 2011), is a key, largely unexplored question that may preclude its support. Other mechanistic models have been proposed (Innan & Kondrashov, 2010), yet do not convincingly resolve the main population genetic dynamical issue.

In this work, we tested the idea of error buffering to reveal the adaptive value that gene duplication has *per se*. Subsequently, we developed a comprehensive model to explain the early fate of duplicates compatible with population genetics (Lynch *et al*., 2001; Lynch, 2007), global gene expression patterns (Qian *et al*., 2010; Gout & Lynch, 2015; Cardoso-Moreira *et al*., 2016; Lan & Pritchard, 2016), and unexpected gene copy number variation rates (Reams *et al*., 2010; Schrider *et al*., 2013). To this end, instead of performing a conventional sequence analysis (top-down approach), we followed a very precise quantitative framework, based on biochemistry, to study the goodness of having a second gene copy for the cell without functional divergence (bottom-up approach). Using a real gene of *Escherichia coli* (*lacZ*) as a model system from which to apply our theory, we showed, without loss of generality, that the sum of two different, partially correlated responses allows reducing gene expression inaccuracies (Rodrigo & Poyatos, 2016); inaccuracies that are a consequence of the inherently stochastic nature of all molecular reactions underlying gene expression (Raser & O’Shea, 2004; Carey *et al*., 2013) and suboptimal gene regulation (Dekel & Alon, 2005; Price *et al*., 2013). In turn, cell fitness can weakly increase on average, if such errors in gene expression are costly (Wang & Zhang, 2011), and genotypes with duplication can then be fixed in the population. We further studied the genetic and environmental conditions that are more favorable for the selection of gene duplication.

## RESULTS

### Quantitative biochemical view of a fitness trade-off

In cellular systems, fitness trade-offs usually arise because beneficial actions involve costs. A paradigmatic trade-off emerges when a given enzyme needs to be expressed to metabolize a given nutrient present in the environment (**Fig. 1a, b, c**). On the one hand, the cell growth rate (here taken as a metric of fitness; Elena & Lenski, 2003) increases as long as the enzyme metabolizes the nutrient. On the other hand, the enzyme expression produces a cost to the cell (*i.e.*, reduces its growth rate). Therefore, the enzyme expression needs to be very precise to warrant an optimal or near-optimal behavior (cost-benefit analysis). To solve this issue, regulations (mainly transcriptional) evolved to link enzyme expression inside the cell with nutrient amount available in the environment. An example of this paradigmatic system is the well-known lactose utilization network of *E. coli* (Jacob & Monod, 1961), where lactose (nutrient, environmental molecule) activates, through inhibition of LacI (transcription factor), the production of LacZ (enzyme). We used this model system to apply a theoretical framework (see Methods) in order to reveal the intrinsic adaptive value of gene duplication under a fitness trade-off, as this system has been quantitatively characterized (Dekel & Alon, 2005; Kuhlman *et al*., 2007; Eames & Kortemme, 2012).

**Figure 1:**
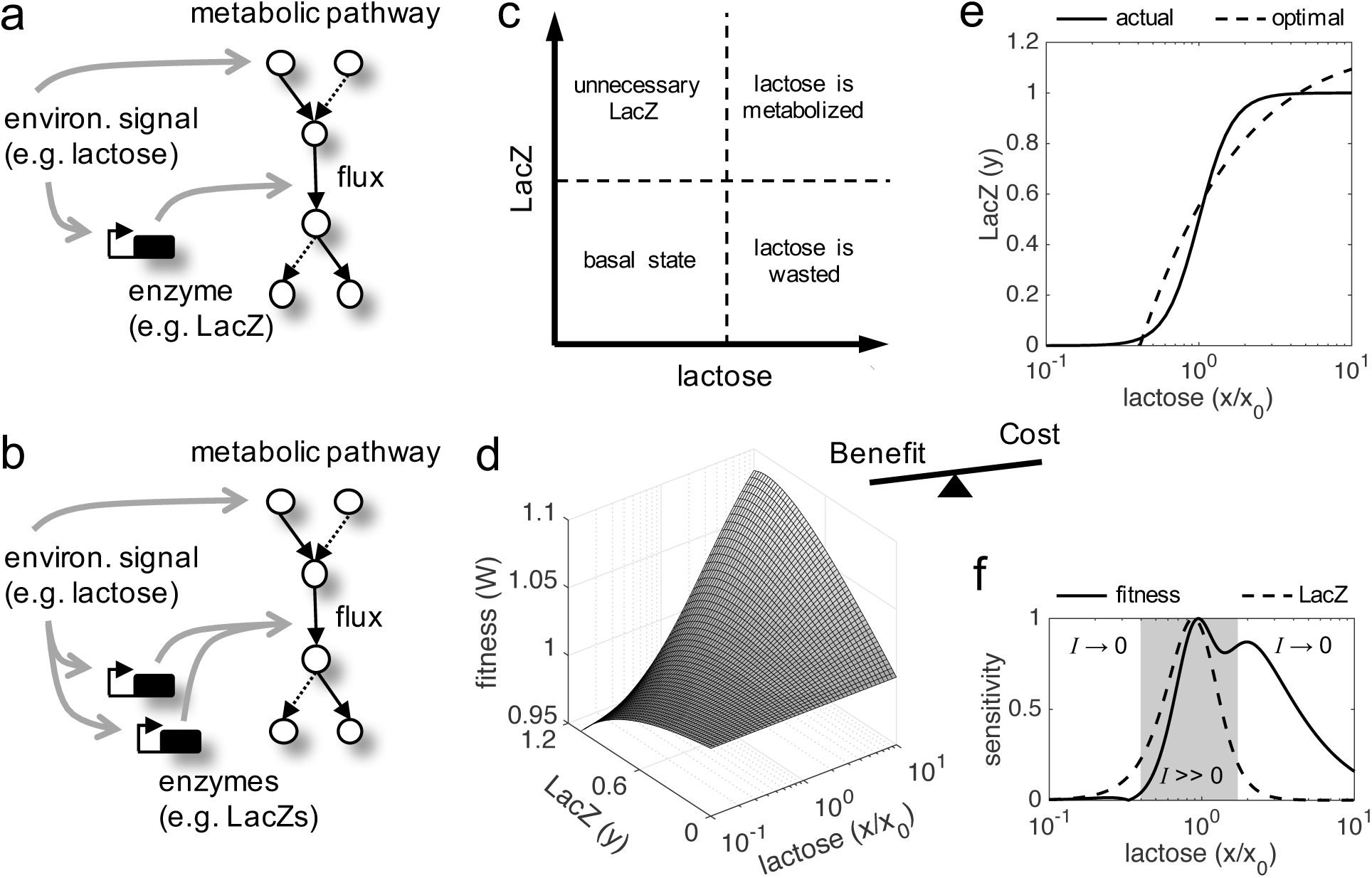
**a)** Scheme of a paradigmatic genetic system, coupling regulation and metabolism, where a given environmental signal determines the physiology of the cell. The environmental molecule can be metabolized by the cell, and can also activate transcriptionally the expression of enzymes. A particular case is the lactose utilization system of *E. coli*. **b)** Scheme of the same system with gene duplication. **c)** Illustrative chart of the fitness trade-off showing four different cellular regimes. When the signal molecule (lactose) is not present in the medium, the expression of the enzyme (LacZ) is not required. However, when the signal molecule is present, the enzyme is required for its metabolic processing. **d)** Fitness (*W*) landscape as a function of lactose (contributing to the benefit, *x* denotes its concentration) and LacZ (contributing to both the benefit and the cost, *y* denotes its concentration). This was experimentally determined. **e)** Dose-response curve between lactose and LacZ. The solid line corresponds to the actual regulation (experimentally determined), whilst the dashed line corresponds to a hypothetical optimal regulation (obtained by imposing *dW* / *dy* = 0). **f)** Sensitivity to changes in lactose dose, either in fitness (*dW* / *dx*, solid line) or in LacZ (*dy* / *dx*, dashed line), characterizing the nonlinear phenotypic plasticity of the cell. Each curve was normalized by its maximum. This also measures sensitivity to molecular noise. The region where information transfer is high is shaded.

Cell fitness increases monotonically with lactose dose, but presents an optimum with LacZ expression (**Fig. 1d**). This is because lactose does not introduce a cost into the system, but LacZ does. Here, we simply considered a cost function based on LacZ expression, although it would be more precise a cost based on lactose permease (LacY) activity (Eames & Kortemme, 2012). The regulation of the system appears to be quite accurate, as the actual and optimal dose-response curves roughly match (**Fig. 1e**). This entails great phenotypic plasticity of the cell to cope with lactose variations. However, plasticity is not equal for all environmental changes. Whilst the system (in terms of LacZ expression or cell fitness) reaches optimal sensitivity at intermediate doses, it is quite insensitive at very low or very high doses, where lactose-LacZ information transfer falls down (**Fig. 1f**).

### Gene duplication helps to better resolve the fitness trade-off

The LacZ expression in *E. coli* involves a variety of noisy actions, such as the LacI expression, the LacI-DNA binding, the RNA polymerase-DNA binding, and the transcriptional elongation process (Elowitz *et al*., 2002; Raser & O’Shea, 2004; Carey *et al*., 2013). Using a simple mathematical model, we simulated the stochastic LacZ expression of the wild-type system for a varying lactose dose (**Fig. 2a, b**). At a given dose, these simulations would correspond to different single-cell responses. We also considered a system with two copies of the *lacZ* gene, with total expression equal to the previous one-copy system, and simulated its stochastic response (**Fig. 2c**). We observed that the system with gene duplication produces a more accurate response (*i.e*., a response closer to the deterministic one), highlighting the role of gene copy number in noise buffering (Rodrigo & Poyatos, 2016). More precisely, we quantified how much information is transmitted through these two systems (1.29 bits of information in case of a singleton and 1.58 bits in case of a duplicate), revealing an increase of about 25% in fidelity when a second gene copy is at play (Hansen & O’Shea, 2015; Rodrigo & Poyatos, 2016). The increase in fidelity is notable as the system presents high regulatory sensitivity (*i.e*., nonlinearity).

**Figure 2:**
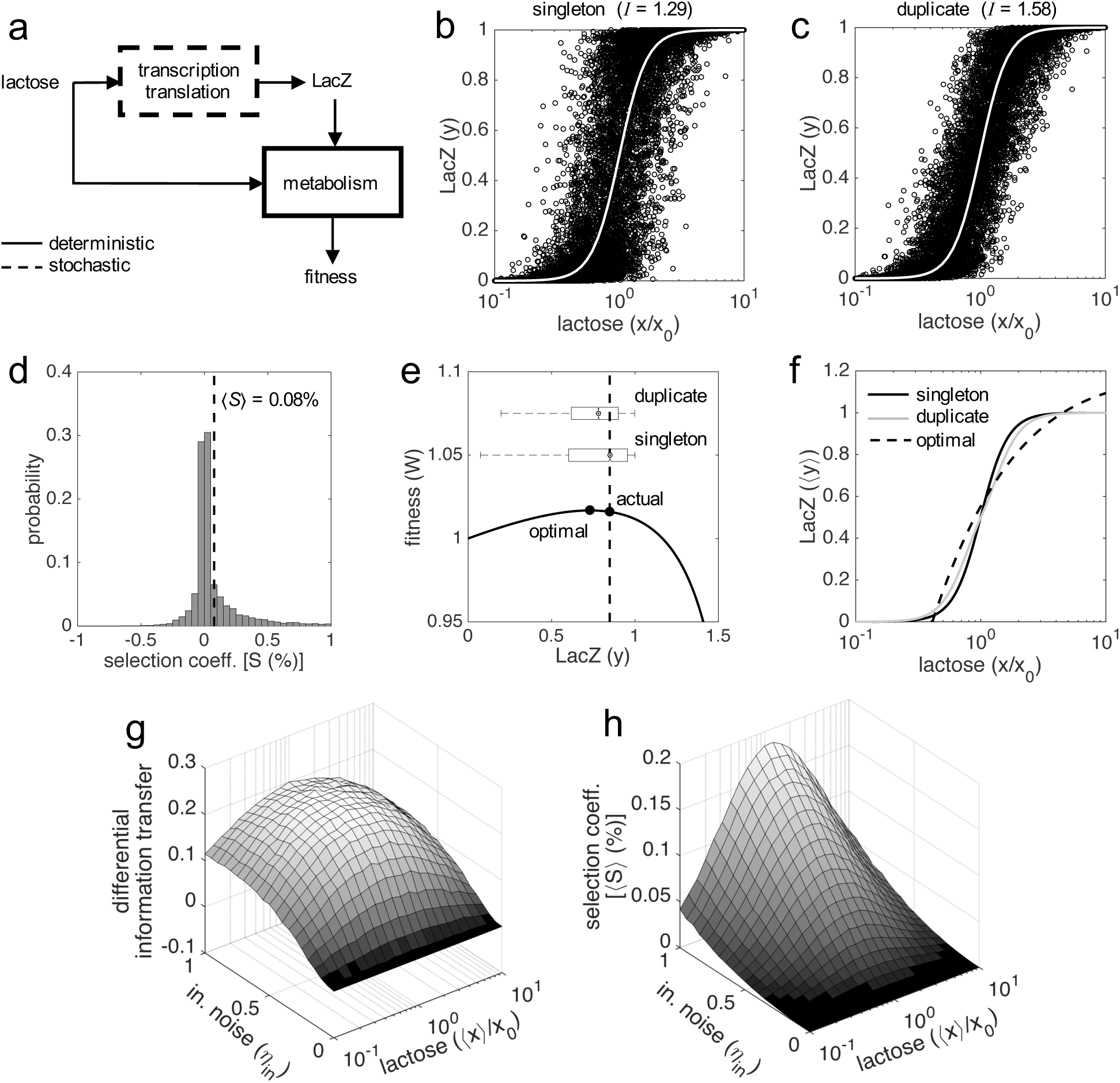
**a)** Block diagram of the system. Gene expression is calculated by means of a stochastic function, whilst fitness by means of a deterministic one. **b, c)** Single-cell responses at different lactose doses (stochastic simulations, noise amplitudes of *η*_in_ = 0.5 and η_ex_ = 0). Lactose and LacZ concentrations are denoted by *x* and *y*, respectively. The solid white line corresponds to the deterministic simulation. In a) the genotype contains a single copy of *lacZ* gene, whilst in b) it contains two copies. The value of mutual information (*I*) is shown in both cases. **d)** Selection coefficient (*S*) of a genotype with two copies of *lacZ* gene over another with just one copy. The mean selection coefficient is shown (dashed line). Skewness coefficient of 2.63. *W* values calculated from *x*, *y* values shown in b, c). **e)** Fitness (*W*) as a function of LacZ (constant *x* = 0.2 mM), showing the distributions of expression (boxplots) in case of one or two gene copies. The actual LacZ expression is shown (dashed line). **f)** Dose-response curve between lactose concentration and the median LacZ expression (〈*y*〉). The solid lines correspond to the actual responses in case of one (black) or two (gray) gene copies, whilst the dashed line corresponds to the optimal response. **g)** Information transfer landscape (measured as the difference in mutual information of duplicate versus singleton) as a function of the median lactose dose (〈*x*〉), fluctuating dose) and the amplitude of intrinsic noise (*η*_in_, with fixed *η*_ex_ = 0.3). **h)** Mean selection coefficient (〈*S*〉) landscape of gene duplication as a function, as in g), of 〈*x*〉 and *η*_in_. In all these plots, the expression levels of the duplicates with respect to the singletons were equal (*y*_max,1_ = *y*_max,2_ = 0.5).

In addition, we calculated the proposed fitness function for each single-cell response. Small gene expression inaccuracies (*e.g*., an excess of enzyme for the available substrate) can be perceived as a consequence of a rugged fitness landscape in terms of the genotype-environment interaction (**Fig. 1d**). To properly compare how each system of study resolves the fitness trade-off, we then calculated the selection coefficient for each response. We found a skewed distribution, peaked at 0 and with a positive mean of 0.08% (**Fig. 2d**). This entails that phenotypic responses generated by duplicated genes give, on average, higher fitness values than responses generated by singleton genes. To better illustrate this fact (which we call phenotypic accuracy), we represented cell fitness as a function of LacZ expression (**Fig. 2e**), uncovering two reasons by which gene duplication is adaptive. In first place, the variance of the stochastic fluctuations in gene expression is reduced by 50% upon duplication (Wang & Zhang, 2011). This increases fitness on average, because the system displays a near-optimal behavior in the deterministic regime, thus fluctuations are costly. In second place, the population response is slightly closer to the optimal operation point (**Fig. 2e, f**). This is because, in this case, the actual dose-response curve is more nonlinear than the optimal one, a feature that can indeed be amended by genetic redundancy (Rodrigo & Poyatos, 2016).

Finally, we calculated how much information is transmitted in the system and how much selection exists, on average, as a two-dimensional function of the magnitude of intrinsic noise and the concentration of lactose in the medium (**Fig. 2g, h**). As expected, the two surfaces resemble, highlighting the fundamental link between phenotypic accuracy and selective advantage (Spearman’s correlation coefficient of 0.927, *P* < 0.001). More in detail, we found that the higher the intrinsic noise, the higher the adaptive value of gene duplication (**Fig. 2h**). This is because intrinsic noise generates the required heterogeneity between the responses of the two gene copies to limit large stochastic fluctuations in the total gene expression. We also found that there is a maximal adaptive value of gene duplication at intermediate lactose doses (**Fig. 2h**), where the sensitivity of the system is the highest (**Fig. 1f**). Out of this regime, the stochastic fluctuations, according to our simple mathematical model, have less impact on the phenotype (Blake *et al*., 2006).

### Gene duplication can be positively selected in a population by phenotypic accuracy

If gene duplication enhances phenotypic accuracy on average, *viz.*, by reducing gene expression inaccuracies, it would be expected a positive selection of this trait in a population (Kimura, 1983). To verify this assumption, we performed experiments of *in silico* evolution (see Methods), where a mixed population of cells carrying singletons and duplicates was monitored, considering equal LacZ expression in both types of cells (**Fig. 3a**). The population was left to evolve without introducing any artifact, with time-dependent stochastic fluctuations in gene expression uncorrelated from cell to cell. We found that the frequency of cells carrying duplicates in the population increases with time, and that such an increase is well predicted by population genetic dynamics with the mean selection coefficient (**Fig. 3b**). Notably, this points out that this parameter, which can be mathematically calculated *a priori*, is sufficient to capture all the complexity underlying the stochastic evolutionary dynamics of the system (Hegreness *et al*., 2006).

**Figure 3:**
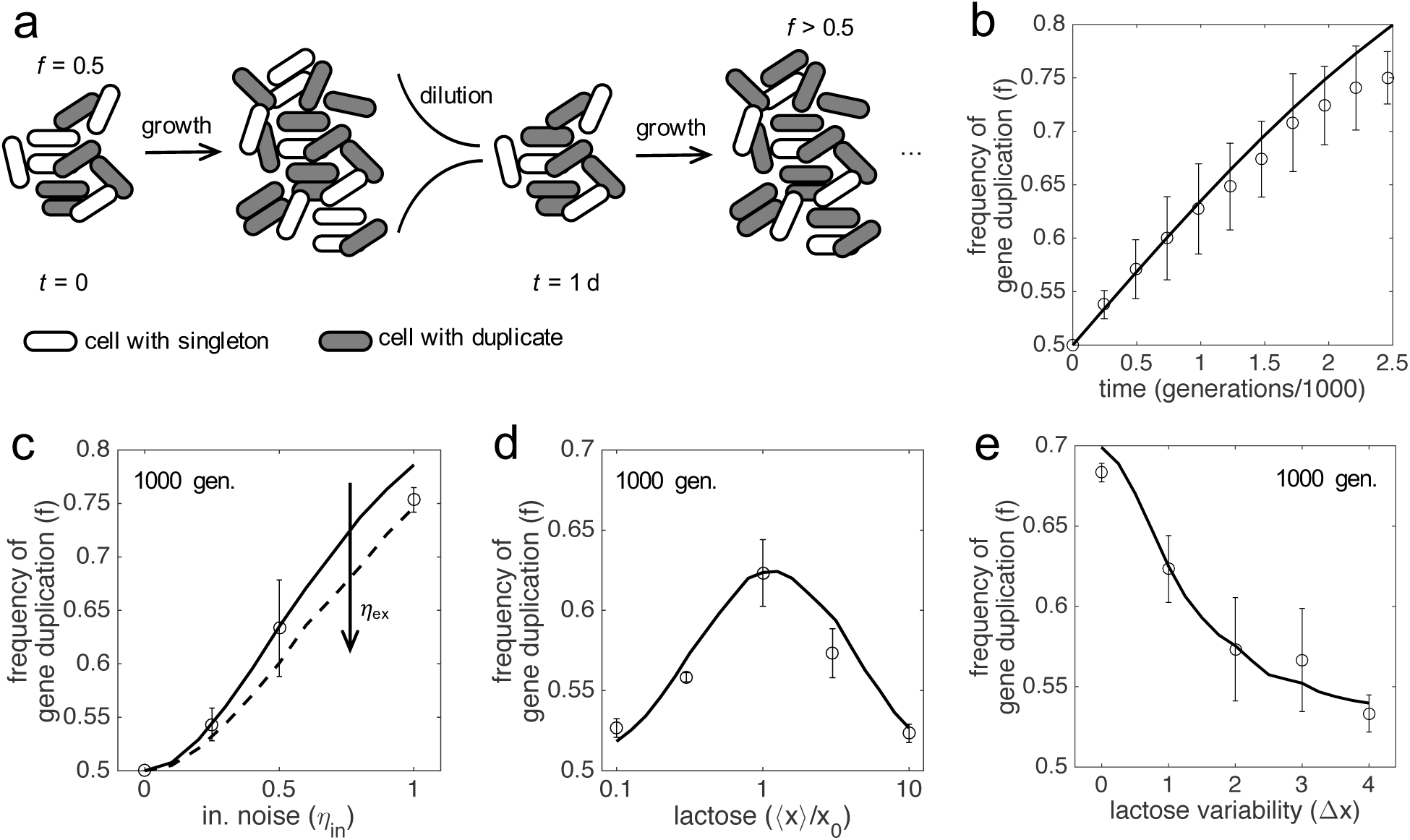
**a)** Scheme of an evolutionary procedure, where serial dilution passages are applied, to assess the performance in a cell population of a genotype with two copies of *lacZ* gene over another with just one copy. **b)** Time-dependent frequency of cells with gene duplication (*f*). Open circles and error bars correspond to experiments of *in silico* evolution (mean and standard deviation of three replicates) with an initial frequency of *f*_0_ = 0.5, fluctuating lactose dose, and noise levels of *η*_in_ = 0.5 and *η*_ex_ = 0. The solid line corresponds to the theoretical prediction. **c)** *f* at 1000 generations (*f*_1000_) as a function of the amplitude of intrinsic noise (*η*_in_). Experiments and prediction with *f*_0_ = 0.5 and *η*_ex_ = 0. The dashed line corresponds to the theoretical prediction with *η*_ex_ = 1. **d)** *f*_1000_ as a function of the median lactose dose (〈*x*〉). Experiments and prediction with *f*_0_ = 0.5, *η*_in_ = 0.5 and *η*_ex_ = 0.5. **e)** *f*_1000_ as a function of the lactose fluctuation amplitude (Δ*x*). Δ*x* = 0 corresponds to constant lactose dose. Experiments and prediction with the same values of *f*_0_, *η*_in_ and *η*_ex_ as in d). Three replicates were also considered in c, d, e). In all these plots, the expression levels of the duplicates with respect to the singletons were equal (*y*_max,1_ = *y*_max,2_ = 0.5).

In addition, we studied the effect of the magnitude of molecular noise. We distinguished between intrinsic and extrinsic noise (Elowitz *et al*., 2002). As predicted from our previous results, we found that the higher the intrinsic noise of the system, the higher the frequency of gene duplication in the population (**Fig. 3c**). By contrast, the higher the extrinsic noise, the lower the frequency (**Fig. 3c**), as this type of noise affects in the same way the responses of the two copies. Note that there is no gain following duplication when only extrinsic noise is considered. Furthermore, we studied the effect of the environment (lactose dose). As predicted, we found an intermediate median dose at which the frequency of gene duplication in the population is the highest (**Fig. 3d**). We also found that the higher the variance, the lower the frequency (**Fig. 3e**). This is because, when lactose fluctuates from very low to very high doses, the signal-to-noise ratio is large enough to warrant a relatively accurate response with just one gene copy (Hansen & O’Shea, 2015). Of relevance, the population genetic dynamics in all these cases, with the corresponding mean selection coefficients, correctly explained the reported frequencies.

### Most of the new-born duplicated genes are costly for the cell and do not offer phenotypic accuracy

So far, we have demonstrated that a duplicated gene offers a selective advantage provided the total gene expression level is maintained, with one or two copies. However, this condition is not met during the critical very initial phase, when the duplicate has just born. In general, we can assume that the expression level is doubled upon duplication, although this may vary due to the particular position in the chromosome of the duplicated gene and the type of cell (Stranger *et al*., 2007). Certainly, an increase of expression due to gene duplication is detrimental in most environments (**Fig. 4a**; Price & Arkin, 2016), thus positive or neutral selective conditions are difficult to invoke to explain the fixation of these type of genotypic changes, mainly in prokaryotes and lower eukaryotes (Lynch & Marinov, 2015). For instance, at constant 0.13 mM lactose, we obtained mean selection coefficients between −28% (very high noise) and −1% (no noise) upon duplication of the *lacZ* gene (assuming double expression), which yield negligible fixation probabilities (almost 0) for a sufficiently large bacterial population. Only in absence of lactose, when the enzyme is not needed, the duplication is strictly neutral (no benefit, no cost due to regulation).

**Figure 4:**
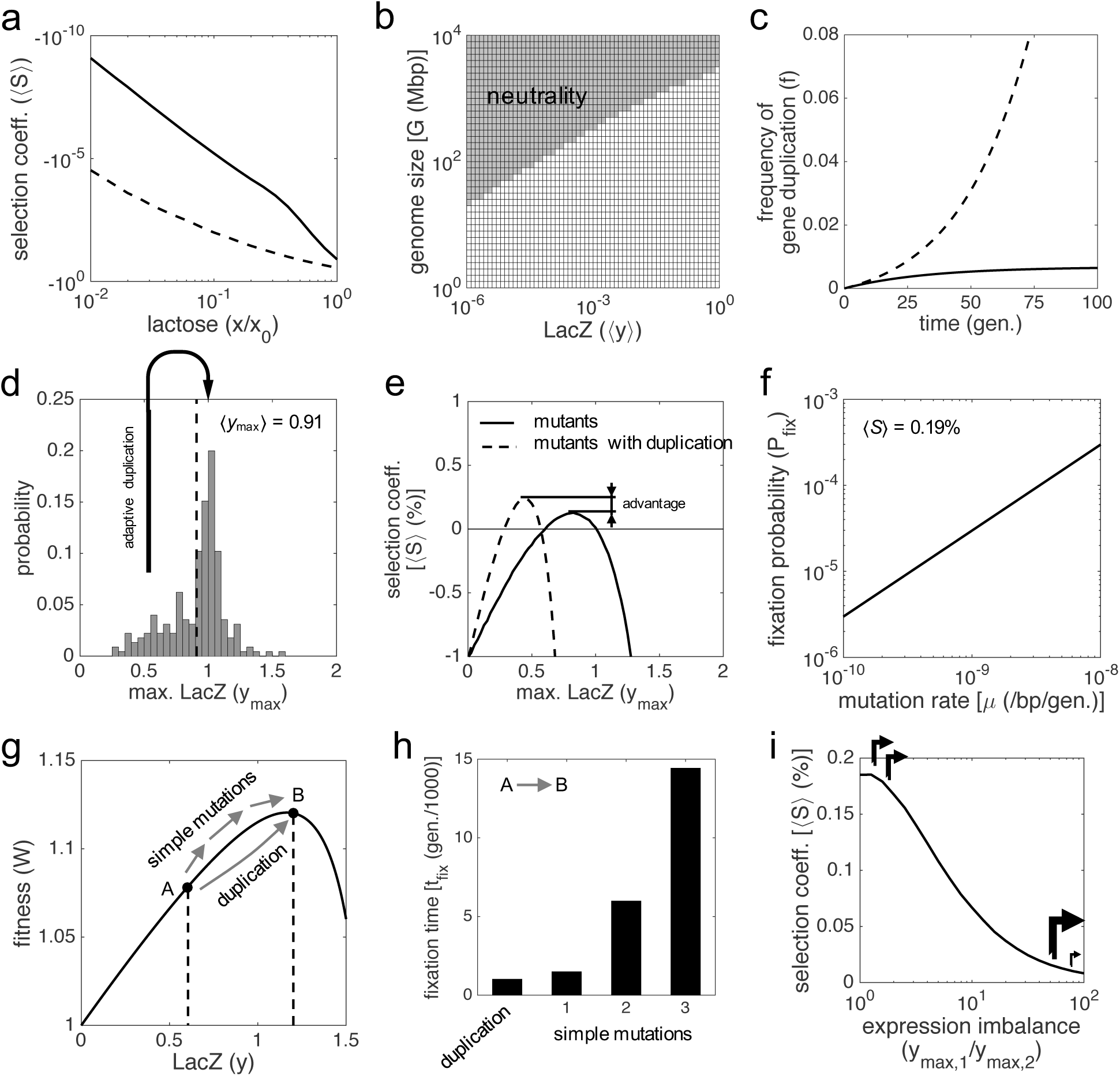
**a)** Mean selection coefficient (〈*S〉*) as a function of lactose dose upon *lacZ* duplication doubling gene expression (*y*_max,1_ = *y*_max,2_ = 1). The solid line corresponds to noise levels of *η*_in_ = *η*_ex_ = 0.3 (moderate), whilst the dashed line corresponds to *η*_in_ = *η*_ex_ = 1 (high). **b)** Identification of effectively neutral selective conditions (when |〈*N*〉·〈*S*〉| < 1, region shaded) in terms of gene expression (*y*) and genome size (*G*), which determines the effective population size (〈*N*〉). In this context, no benefit was considered (*a* = 0), with moderate noise levels. **c)** Time-dependent frequency of cells with gene duplication (*f*) when the creation and deletion rates of a second *lacZ* copy are considered. Sequence remodeling was not taken into account. The solid line corresponds to a scenario of neutrality, whilst the dashed line corresponds to a scenario of positive selection (with *S* = 10%). **d)** Distribution of the activity of *lac* promoter mutants based on experimental data, as the maximal LacZ expression (*y*max, irrespective of lactose dose). The mean activity is shown (dashed line). Skewness coefficient of −0.68. **e)** 〈*S*〉 of the promoter mutants versus the wild-type system (solid line), with fluctuating lactose dose, and high noise levels. The dashed line corresponds to the comparative between promoter mutants that duplicated the *lacZ* gene and the wild-type system. **f)** Fixation probability (*P*_fix_) of gene duplication as a function of the mutation rate of the cell (p), with 〈*S*〉 = 0.19%. **g)** Fitness (*W*) as a function of LacZ (constant *x* = 6.5 mM), with *y*_max_ = 0.6. Point A designates the ancestral genotype, whilst point B designates a new genotype with double LacZ expression generated by gene duplication or by simple mutations. **h)** Fixation times (*t*_fix_) of different adaptive genotypes, with 〈*N*〉 = 10^9^, and moderate noise levels. In case of duplication or one simple mutation that doubles gene expression (μ_b_ = μ), the mean selection coefficient is 〈*S*〉 = 3.9%. In case of two successive mutations with equal increase in expression (μ_b_ = 3·μ), the coefficients are 〈*S*〉 = 2.79% for the first one, and 〈*S*〉 = 1.09% for the second one. In case of three successive mutations with equal increase in expression (μ_b_ = 10·μ), the coefficients are 〈*S*〉 = 1.97% for the first one, 〈*S*〉 = 1.44% for the second one, and 〈*S*〉 = 0.45% for the third one. **i)** 〈*S*〉 as a function of the expression imbalance between the two *lacZ* copies (*y*_max,1_ / *y*_max,2_), when the system recovers its ancestral expression levels (*y*_max,1_ = *y*_max,2_ = 0.5), with constant *x* = 0.13 mM, and high noise levels.

Even though, neutral selective conditions can be reached *de facto* if the absolute value of the selection coefficient is lower than the inverse of the effective population size (Kimura, 1983). This condition is challenging for prokaryotes, as their population sizes are very large (Lynch & Conery, 2003). In our particular case, we obtained mean selection coefficients in the order of −10^−10^ (moderate noise) when the nutrient amount is scarce (1 μM lactose), which could favor the fixation of a *lacZ* duplicate by genetic drift. It can be argued, nevertheless, that the cost of over-expression decreases as long as the genome size increases (Lynch & Marinov, 2015). This assumption, together with the negative correlation between complexity and population size (Lynch & Conery, 2003), makes effectively neutral selective conditions plausible to rationalize the fixation of duplicates that are expressed (*e.g*., essential genes) in higher eukaryotes (**Fig. 4b**; Makino *et al*., 2009).

### Fixation is conditioned by the unexpected recurrence of creation and deletion of gene duplications in a population

Gene duplications can be spontaneously created, through different mechanisms (Hastings *et al.*, 2009), at very high rates in the cell. These rates, measured from experiments of mutation accumulation, go from 10^−4^ dup./gene/gen. in prokaryotes (Reams *et al*., 2010) to 10^−7^ dup./gene/gen. in higher eukaryotes (Schrider *et al*., 2013). Once created, most of these duplications are deleted as they are unstable, with a rate that appears to be higher than the creation rate (Reams *et al*., 2010; Schrider *et al*., 2013). In the particular case of the *lacZ* gene, we have a creation rate of 3·10^−4^ dup./gene/gen. and a deletion rate of 4.4·10^−2^ -/gene/gen. (in a single bacterial cell). Therefore, gene duplication can be understood as a recurrent process that reaches an equilibrium point given by the ratio between the creation and deletion rates, neglecting fitness effects. This equilibrium point would be lower if fitness effects (mostly detrimental) were considered. This entails about 10^6^ cells carrying *lacZ* duplicates in a typical bacterial population of 10^9^ cells (*i.e.*, frequency of about 0.1%). This surprising scenario has an immediate consequence, *viz.*, duplicated genes cannot be fixed in the population by drift under neutral selective conditions (**Fig. 4c**); a result already anticipated (Clark, 1994) in clear discrepancy with the conventional wisdom (Lynch, 2007). Indeed, the creation-deletion balance would always take the system to the same equilibrium point.

However, the preceding argument only focuses on a static picture, ignoring the dynamics of the genetic process. In bacteria (*lacZ* gene), the time to reach the equilibrium point is about 68 generations (three times the inverse of the deletion rate), which is a relatively short transient period. By contrast, in flies (*Drosophila melanogaster*), the creation rate is of 10^−7^ dup./gene/gen. and the deletion rate of 10^−6^ -/gene/gen. (Schrider *et al*., 2013). Although this would yield equilibrium frequencies up to 10%, the transient periods would be longer than 10^6^ generations (0.2 Ma in natural conditions; Pool, 2015). Fixation could then happen by drift, as their effective population sizes are of 10^6^ flies, although not persistently. Note that the inverse of this number indeed specifies an upper limit for the deletion rate. In addition, the creation-deletion balance could be shifted if further genome rearrangements affecting duplicated genes were considered, such as gene relocation (about 10^−11^ fixed rearr./gene/gen. for *D. melanogaster*; Ranz *et al*., 2001). In effective terms, gene relocation would reduce the deletion rate, and, consequently, fixation would be more likely (Wong & Wolfe, 2005). Such a relocation would also shift the intrinsic-extrinsic noise balance towards more uncoupled responses (Becskei *et al*., 2005), which could enhance phenotypic accuracy.

### Phenotypic accuracy can lead to the fixation of a new-born duplicated gene in the population

Can a cell carrying a new-born duplicate that is expressed overcome the additional cost and then invade the population by phenotypic accuracy? We here predicted that the genetic variability existing in a population would allow reaching adaptive gene duplications. Mutations in the *cis*-regulatory region of the *lacZ* gene may change its wild-type expression level. According to previous results (Otwinowski & Nemenman, 2013), the distribution of mutations in terms of maximal promoter activity is peaked at 1, but skewed to the left (**Fig. 4d**). This indicates that most of these mutations are nearly neutral, but some of them (about 10%) yield cells with nearby 50% lower expression. Thus, if a gene duplication event occurred in one of these cells, the genotypic change would be selectively advantageous (**Fig. 4e**). The frequency of such cells in the population depends, of course, on the mutation rate; the greater the ability to generate genetic diversity, the higher the chances to reach adaptive duplications. For *E. coli*, where the simple mutation rate is of 10^−10^ mut./bp/gen. (Lee *et al*., 2012), this frequency can be estimated in 10^−9^ (*i.e*., 1 mutant with nearby 50% lower expression in a population of 10^9^ cells). Hence, the probability that a duplication and such a mutation concur in the same cell is of 10^−3^ (*i.e.*, 1 concurrence each 10^3^ generations).

In particular, at constant 0.13 mM lactose, we obtained a relatively high mean selection coefficient of 0.19% when the wild-type expression is recovered upon duplication (in a highly noisy scenario). However, the selection coefficient has to be greater than the duplication deletion rate to ensure fixation (**Fig. 4c**); a condition that is not met here. Certainly, the high deletion rates observed in bacteria (Reams *et al*., 2010) protect them from acquiring genetic redundancy. In other local genetic contexts, also in bacteria, the deletion rate of a *lacZ* duplicate can be as low as 4.1·10^−4^ -/gene/gen. (Reams *et al.*, 2012). In this scenario, a selection coefficient of 0.19% would lead to fixation. We then estimated a global fixation probability of 3·10^−6^ (**Fig. 4f**; see Methods). Remarkably, our estimation is much higher than 10^−9^, the fixation probability under hypothetical neutrality (Kimura, 1983).

Phenotypic accuracy could also lead to the fixation of duplicates in eukaryotes, as nothing prevents assuming the same positive selective conditions (Raser & O’Shea, 2004; Hansen & O’Shea, 2015), which now largely outperform the duplication deletion processes. For *D. melanogaster*, for instance, where the simple mutation rate is of 5·10^−9^ mut./bp/gen. (Schrider *et al*., 2013), and complete gene duplications have little impact on fitness (Emerson *et al*., 2008; note that other genome rearrangements not affecting entire genes are significantly deleterious), we estimated that 0.05 mutants with nearby 50% lower expression, and 10^5^ duplicants of the gene of interest would be found in the natural population. Hence, the probability of concurrence in the same organism would be of 5·10^−3^. Consequently, the global fixation probability would be of 2·10^−5^; again, higher than the one under hypothetical neutrality (Kimura, 1983).

### Expression demand in extreme environments can also lead to the fixation of a newborn duplicated gene in the population

Gene duplications can be adaptive in response to environments that demand more expression. In general, these are extreme environments (Riehle *et al*., 2001). In the case of the lactose utilization network, very high nutrient amounts require considerable enzyme sums to optimize the cell growth rate. We now considered, for illustrative purposes, a *lac* promoter with 40% lower activity. The expression levels achieved by this system are still acceptable to cope with lactose doses lower than 0.13 mM, but not to cope with very high doses (extreme environments). Indeed, the system would require double LacZ expression at constant 6.5 mM lactose (**Fig. 4g**). In this regard, two different adaptive trajectories can happen. On the one hand, a gene duplication event occurs and the resulting genotype fixes in the population. Note that the mean selection coefficient is, in this scenario, much higher (3.90%) than previously shown (**Fig. 2h**). On the other hand, simple mutations that increase the *lac* promoter activity occur and the resulting genotype, with one or more mutations, also fixes in the population.

Hence, this generates a competition between the duplicate and the simple mutants to face the extreme environment, a phenomenon known as clonal interference in asexual populations (Rozen *et al*., 2002; Desai *et al*., 2007). The fixation time of a duplication occurring immediately can be estimated in 10^3^ generations for *E. coli*, which is compatible with previous experimental results (Riehle *et al*., 2001), and it is similar to the fixation time of one simple mutation that doubles gene expression (**Fig. 4h**; considering serial dilution passages). Note also that the selective advantage is equal in both cases (*i.e*., there is no gain in accuracy in case of duplication), as the system is in a region of poor information transfer (**Fig. 1f**). In addition, the fixation time of genotypes that require two or more mutations is higher (**Fig. 4h**). Accordingly, we predicted that if just one mutation already optimizes the expression, duplications will be fixed only in about half of the cases. However, when two or more mutations are needed, the fixation of the duplicate will be more likely. In turn, if the expressions of multiple genes (*e.g*., a gene cluster) were needed to be increased for adaptation, a duplication of the entire cluster would be an efficient solution (Pollard & Holland, 2000; Riehle *et al*., 2001).

Expression demands in extreme environments could also dictate the fixation of duplicates in complex organisms. In this case, the fixation time (in absolute terms) would be much longer, as these organisms reproduce at larger time scales (weeks or years); in contraposition to doubling times of minutes or hours in simple organisms. Therefore, the extremality would have to remain for very long time to allow fixation in the population (step-like). For instance, the continued use of insecticides has shaped the genomes of flies by promoting the fixation of duplications of toxin-resistance genes (Emerson *et al*., 2008). In this regard, complex organisms could not face a punctually appearing, shortly ceasing extremality (pulse-like). However, if the frequency of such environments were significant, they could develop bet-hedging strategies (King & Masel, 2007).

### Initial maintenance upon fixation in the population of a duplicated gene

A forthcoming change in lactose dose would be highly detrimental if a second *lacZ* copy were fixed in the population either under neutrality due to insignificant expression, or under strong selection due to expression demand. In the former case, an increase would be detrimental; in the latter, a decrease would. Consequently, either the elimination of the duplicate by purifying selection (Lynch & Conery, 2000), or the accumulation of mutations that lower the LacZ expression to recover the ancestral phenotype (Force *et al*., 1999; Qian *et al*., 2010) would be promoted. In the latter case, the two gene copies could be maintained in the genome for long time by phenotypic accuracy if they held similar expression levels (**Fig. 4i**; Gout & Lynch, 2015); otherwise the gain in accuracy decreased. Conversely, if a second *lacZ* copy were fixed by phenotypic accuracy, it would be safe from changes in lactose dose.

### A comprehensive model compatible with population genetics to explain the early fate of gene duplications

Taking all our results together, we formulated a comprehensive model to explain the early fate (*viz*., fixation or elimination) of gene duplications (**Fig. 5**). Notably, this model is compatible with population genetics, involving positive and neutral selective conditions (Lynch, 2007). On the one hand, a significant number of duplicates could be fixed by genetic drift only in complex organisms (*i.e*., higher eukaryotes; sector A in **Fig. 5**). This would be due to their increased ability to allocate additional resources for expression (Lynch & Marinov, 2015), and their apparently reduced duplication deletion rate with respect to the inverse of the population size (Schrider *et al*., 2013). However, these fixed duplications would not be stable, due to the creation-deletion balance (Reams *et al*., 2010), and then, for a long-term preservation, they would require the accumulation of beneficial mutations (Han *et al*., 2009), or the relocation of the second copy in the genome to prevent its deletion (Ranz *et al*., 2001). This would lead to late fates of sub- or neofunctionalization (Force *et al*., 1999; Conant & Wolfe, 2008).

**Figure 5:**
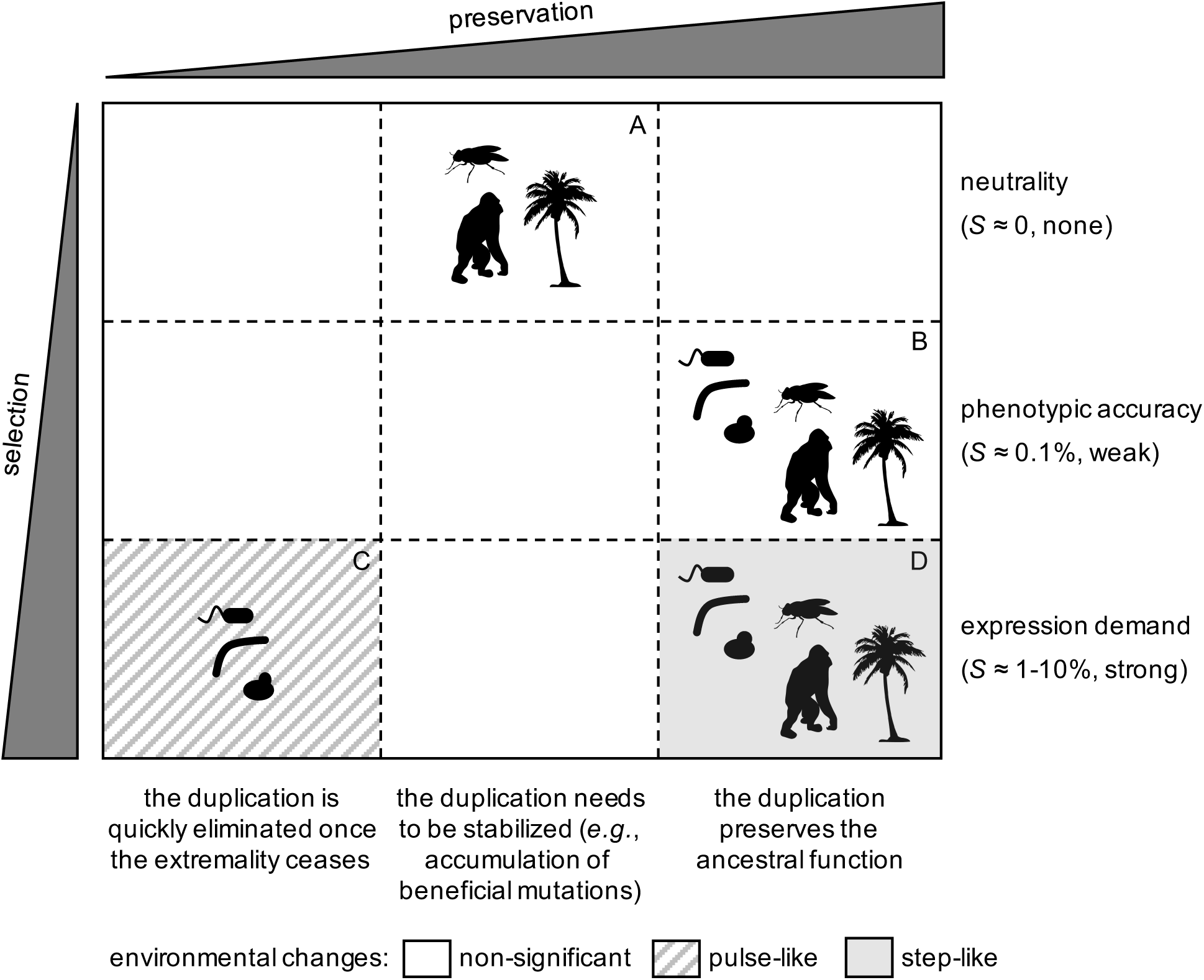
General model to explain the fixation of duplicated genes as a function of the degree of selection in the population, and preservation in the genome for long time. Representative silhouettes correspond to bacteria (prokaryotes), yeasts (lower eukaryotes), insects, plants, and mammals (higher eukaryotes).

On the other hand, positive selection could drive the fixation of duplicates in both complex and simple organisms. When the environmental changes were relatively rapid, only simple organisms (*i.e*., prokaryotes and lower eukaryotes) could fix duplications, due to their short generation times (sector C in **Fig. 5**; Riehle *et al*., 2001). However, such duplications would be quickly eliminated from the population afterwards (once the environment changed again), as the genome rearrangement rates are orders of magnitude higher than the simple mutation rates (Reams *et al*., 2010). By contrast, when a given environmental change were prolonged, complex organisms could also fix duplications (sector D in **Fig. 5**; Emerson *et al*., 2008). In this case, they would be under strong positive selection, and, consequently, they would be preserved for long time. Furthermore, all organisms could fix duplications by phenotypic accuracy (sector B in **Fig. 5**), without the need of significant environmental changes; provided the gene of interest were noisily expressed (Elowitz *et al*., 2002; Raser & O’Shea, 2004), and the duplication deletion rate were lower than the weak selective advantage. Thus, we predicted that a significant number of duplicates observed in the sequenced genomes, especially in simple organisms, would have been fixed by this mechanism. In the very long term, these weak selective conditions could also allow the exploration of novel functions, as they ensure the preservation of duplicates, without invoking fortuitous exploration in the ancestral state (Bergthorsson *et al*., 2007), and with amplification when the advantage provided by the narrowed novel function were higher than the advantage by phenotypic accuracy.

## DISCUSSION

The inherently stochastic nature of gene expression is certainly an evolutionary driver when it is linked to cell fitness to dictate the selection of particular genetic architectures (Batada & Hurst, 2007; Maamar *et al*., 2007). Our results demonstrate that gene duplication can be positively selected as an architecture that allows enhancing information transfer in genetic networks (*i.e*., mitigation of expression errors; Rodrigo & Poyatos, 2016); a feature that we call phenotypic accuracy. This entails that the genetic robustness indeed observed upon the accumulation of genetic redundancy (Keane *et al*., 2014) would be more a consequence than a selective trait (Kafri *et al*., 2006). Moreover, our results highlight that a population genetic model with the mean selection coefficient is enough to explain the complex, stochastic evolutionary dynamics of duplication fixation. Of note, the reported intrinsic adaptive value, which cannot be captured by sequence analyses, was derived from basic mathematical models of gene regulation and cell fitness (Dekel & Alon, 2005).

Notably, we anticipated a series of testable results by following our theory. First, most of the duplicates that appear in the genomes would be under strong purifying selection (Lynch & Conery, 2000). Hence, the total gene expression of those that became fixed is expected to be similar to the expression of the corresponding singletons (*i.e*., gene dosage sharing), with the aim of minimizing deleterious fitness effects. This would hold for many duplicates in both simple and complex organisms (Qian *et al*., 2010; Gout & Lynch, 2015; Cardoso-Moreira *et al*., 2016; Lan & Pritchard, 2016), then preserving the ancestral function (DeLuna, 2008). It is also expected, only in case of complex organisms, the existence of fixed duplicates with increased gene expression levels (Cardoso-Moreira *et al*., 2016), which would reflect the effect of genetic drift (Lynch & Conery, 2003) and also, to a lesser extent, of positive selection after prolonged environmental changes (*e.g.*, the case of flies; Emerson *et al*., 2008).

Second, noisy genes are expected to be more duplicable, especially in simple organisms (*e.g.*, yeast; Dong *et al*., 2011). Indeed, the gain experimented by the system upon duplication is greater when gene expression inaccuracies are significant (Rodrigo & Poyatos, 2016). This would explain the TATA box enrichment in the *cis*-regulatory regions of duplicated genes, as these genetic motifs are associated to high plasticity (*i.e*., high sensitivity to multiple environmental changes) and high gene expression noise by inducing transcriptional bursts (Blake *et al*., 2006; Lehner, 2010). Moreover, essential genes are expected to be less duplicable (He & Zhang, 2006) as a consequence of their reduced gene expression noise (Batada & Hurst, 2007). Genes under the control of regulatory structures that buffer noise (*e.g.*, negative feedbacks) would not be duplicable either (Warnecke *et al*., 2009). In more complex organisms (*e.g.*, mammals), however, a significant fraction of duplicates would accumulate beneficial mutations (Han *et al*., 2009) to ensure preservation for long time, resulting in genes that would be useful or even essential (Makino *et al*., 2009).

Third, the local genetic context would be highly determinant of the fixation of a duplicate (Reams *et al.*, 2012), explaining why some genes are more duplicable than others in scenarios of apparent neutrality (hot spots; Perry *et al*., 2006). Moreover, duplicates would be much shorter lived in prokaryotes than in eukaryotes (Lynch & Conery, 2003). After all, the precise experimental determination of the molecular rates of gene copy number variation would unveil to what extent natural selection has actually rivaled random genetic drift to shape complexity along the course of life history.

## METHODS

### Fitness function

The *lac* operon of *E. coli* (Jacob & Monod, 1961) was considered as a biological model system from which to apply a mathematical framework, and cell growth rate was taken as a metric of fitness (*W*; Elena & Lenski, 2003). In this particular case, the benefit function reads *B* = *a*·*y*·*x* / (*k* + *x*), where *a* accounts for the increase in growth rate due to lactose utilization (*x* denotes its concentration; *y* denotes the normalized LacZ expression), and *k* is the Michaelis-Menten constant. In addition, the cost function reads *C* = *b*·*y* / (*h* - *y*), where *b* accounts for the decrease in growth rate due to LacZ expression, and *h* for the maximal resources available in the cell (Dekel & Alon, 2005). Thus, the fitness function reads *W* = *W*_0_·(1 + *B* - *C*), where *W*0 is the cell growth rate in absence of lactose (*x* = 0). Note that this model underestimates the adaptive ability of the bacterium by not considering the effect of LacY. Moreover, the normalized LacZ expression, in the deterministic regime, is given by *y* = *x^n^* / (*x*_0_*^n^* + *x^n^*), where *x*_0_ is the lactose regulatory constant, and *n* the Hill coefficient (accounting for the regulatory sensitivity). In this model, LacZ is not expressed in absence of lactose. If *y* > *h*, we assumed *W* = 0. All parameter values were experimentally fitted, resulting in *W*_0_ = 1 h^−1^, *a* = 0.17, *k* = 0.40 mM, *b* = 0.036, *h* = 1.80, *x*_0_ = 0.13 mM, and *n* = 4 (Dekel & Alon, 2005). The optimal LacZ expression (*y*_opt_) was obtained by imposing *dW* / *dy* = 0, resulting in *y*_opt_ = *h* - [*b*·*h*·(*k* + *x*) / (*a*·*x*)]^1/2^.

### Stochastic gene expression

The normalized LacZ expression in presence of molecular noise was modeled, in steady state, as *y* = *y*_max_· (*x*·*z*_1_ ·*z*_0_)^*n*^ / [*x*_0_*^n^* + (*x*·*z*_1_ ·*z*_0_)*^n^*], where *y*_max_ is the maximal expression level (in general, *y*_max_ = 1), and *z*_1_ and *z*_0_ random variables accounting for intrinsic and extrinsic noise sources, respectively. Here, they were log-normally distributed [with mean 0 for both log(*z*_1_) and log(*z*_0_), and standard deviation *η*_in_ for log(*z*_1_) and *η*_ex_ for log(*z*_0_)]. This accounts for the noisy de-repression of the promoter and subsequent expression due to lactose. Note that whilst LacZ can show a bistable expression pattern with non-metabolizable synthetic compounds (Ozbudak *et al*., 2004), its expression is monostable with lactose (van Hoek & Hogeweg, 2006). For simplicity, the transient LacZ expression was overlooked, and the noise levels were considered constant during a cell cycle. The median response of a population is denoted by 〈*y*〉.

### Gene duplication

The combined expression of two genes coding for LacZ in presence of molecular noise was modeled as *y* = *y*_max,1_·(*x*·*z*_1_·*z*_0_)^*n*^ / [*x*_0_*^n^* + (*x*·*z*_1_·*z*_0_)*^n^*] + *y*_max,2_·(*x*·*z*_2_·*z*_0_)^*n*^ / [*x*_0_*^n^* + (*x*·*z*_2_·*z*_0_)*^n^*], where *z*_2_ is a random variable accounting for intrinsic noise on the second copy, with the same distribution as for *z*_1_ (*z*_1_ and *z*_0_ as before). Note that whilst extrinsic fluctuations (*z*_0_) are common, intrinsic fluctuations (*z*_1_ and *z*_2_) are independent for each gene copy (Elowitz *et al*., 2002). Moreover, the expression levels of the duplicates with respect to the singletons can be adjusted with the values of *y*_max,1_ and *y*_max,2_, with *y*_max,1_ = *y*_max,2_ = 0.5 for equal total expression, and *y*_max,1_ = *y*_max,2_ = 1 for double expression.

In addition, the bacterial model was modified to simulate the effect of gene duplication in organisms of different complexity. For that, the parameter *h* in the cost function was set in terms of the genome size (*G*, in Mbp of haploid genome), simply as *h* ≈ 0.36·*G* (*e.g*., *G* ≈ 5 for *E. coli*, or *G* ≈ 3000 for *H. sapiens*), assuming that complex organisms have more resources to accommodate new gene expressions (Lynch & Marinov, 2015). The effective population size (here denoted by 〈N〉), determinant of the fixation of new genotypes, was also set in terms of *G*, resulting in 〈*N*〉 ≈ 3·10^9^ / *G*^1.44^; an equation roughly inferred from previously reported estimates (Lynch & Conery, 2003).

### Information transfer

Mutual information (*I*) was used as a metric to characterize information transfer by considering the system as a communication channel between the environmental molecule (lactose) and the functional protein (enzyme, LacZ) resulting from gene expression. *I* was calculated as previously done (Rodrigo & Poyatos, 2016), between log(*x*) and *y*. To model the variation of lactose, a random variable log-normally distributed was considered [with mean 0 and standard deviation 1, otherwise specified, for log(*x* / *x*_0_)]. The median lactose dose is denoted by 〈*X*〉, and the fluctuation amplitude, denoted by Δ*x*, corresponds to the standard deviation of log(*x*).

### Genotype-phenotype map

Here, the LacZ expression defines the phenotype of the cell (*i.e*., its metabolic capacity), and for the wild-type genotype it is lactose dependent through the LacI regulation (Jacob & Monod, 1961). Because differences in fitness are very small, the normalized expression (*y*) was assumed independent of it (Klumpp *et al*., 2009). Potential beneficial mutations are those that change the *lac* promoter activity (the *cis*-regulatory regulatory region of LacZ, of about 10^2^ bp). According to an analysis of a large library of mutants (Kinney *et al*., 2010) resulting in a linear model of categorical variables (Otwinowski & Nemenman, 2013), the distribution of maximal LacZ expression upon single-point mutations was inferred. For simplicity, no epistatic interactions were taken into account, although they could matter.

### *In silico* evolution

A medium with maximal capacity for *N* = 10^5^ cells was considered, and serial dilution passages were simulated (Elena & Lenski, 2003), with a dilution factor of *D* = 100 (in terms of volume, with deterministic dominance). The dilution period was set to 1 d. Lactose also varied with the same period. The doubling time of a given cell was 1/*W*, with *W* calculated from the stochastic LacZ expression. In case of no saturation, the cell volume increased as 2*^W·t^*, where *t* is the time in h. Because doublings occur in about 1 h, the number of generations per passage is bounded to log_2_(*D*) = 6.64. Two genotypes were put in competition: one with a single copy of LacZ, the other with two copies. No mutations were allowed to occur.

### Population genetics

In scenarios of competition between two subpopulations (*i.e*., two different genotypes), the ratio between them (*r*) reads *r* = *r*_0_·2*^S^*^·*t*^, where *r*_0_ is the initial ratio, *S* the selection coefficient, and *t* the time measured in generations (Hegreness *et al*., 2006). By setting *W* and *W’* the fitness values of each genotype (with *W*’ > *W*), the selection coefficient is calculated as *S* = *W*’ / *W* - 1. When fitness changes over time, the mean selection coefficient (〈*s*〉) is used. The frequency of the genotype with advantage in the population is *f* = 1/(1 + 1/*r*). The dynamics of a punctual beneficial mutant appeared in an evolutionary experiment of serial dilution passages, with maximal population size *N* and dilution factor *D*, is given by *r* = 2^*S t*^ / 〈*N*〉, where 〈*N*〉 = *N* / *D*^1/2^ is the geometric mean population size (also considered the effective population size; Lewontin & Cohen, 1969). The fixation probability is *P*_fix_ = 2*S*, and the characteristic fixation time *t*_fix_ = log_2_(〈*N*〉^2^) / *S*. Note that the time for 50% invasion of the population is *t*_half-fix_ = log_2_(〈*N*〉) / *S* = *t*_fix_ / 2. However, we have *P*_fix_ = 1/〈*N*〉 and *t*_fix_ = 2〈*N*〉 for a neutral mutant (Kimura, 1983).

By contrast, if multiple beneficial mutants are recurrently created at rate μ_b_, the dynamics is given by *r* = μ_b_·*N*·2*^S^*^·*t*^ / [*S*·log(*D*)·〈*N*〉] ≈ μ_b_·2*^S^*^·*t*^ / *S*, as in each passage μ_b_·*N* different mutants are generated (valid for μ_b_·*N* > 1; Desai *et al*., 2007). Because mutants are now recurrent, *P*_fix_ = 1, and the characteristic fixation time reads *t*_fix_ = log_2_[〈*N*〉·*S* / μ_b_] / *S*. When *m* different mutations accumulate successively, *t*_fix_ ≈ *t*_fix_(*m*) + *t*_half-fix_(*m*-1) + … + *t*_half-fix_(1), *i.e*., a subsequent mutation can start its fixation when the preceding mutation has invaded the 50% of the population (Lang *et al*., 2013). If μ_b_·*N* << 1, the system can be treated as in the case of a punctual beneficial mutation, and the dynamics can be written as *r* = 2^*S*·(*t* - *T*)^ / 〈*N*〉, with a delay of *T* = log_2_(*D*) / (μ_b_·*N*), the mean number of generations required to create a mutant, and *P*_fix_ = 2*S.*

Moreover, in case of gene duplication, if multiple beneficial mutants are recurrently created at rate μ_c_, and deleted at rate μ_d_, the dynamics is given by *r* ≈ μ_c_·2*^S’^*^·*t*^ / *S’*, with *S’*= *S* - μ_d_ as an effective selection coefficient (valid for μ_c_·*N* > 1, and *S* > μ_d_). Again, if μ_c_·*N* << 1, the system can be treated as in the case of a punctual beneficial mutation, with *P*_fix_ = 2*S’*. If *S* << μ_d_, the stationary solution can be approached by *r* ~ μ_c_ / μ_d_ for effectively neutral mutations, or by *r* ≈ μ_c_ / (μ_d_ - *S*) for deleterious mutations.

### Genetic diversity

The simple mutation rate of *E.coli* is μ = 10^−10^ mut./bp/gen. (Lee *et al*., 2012). Cultures of this bacterium may reach population sizes up to *N* = 10^10^ cells (〈*N*〉 = 10^9^). This means, on average, 0.1 (= μ·〈*N*〉) mutants of a given base pair in the population. The number of base pairs, mainly in the *cis*-regulatory regulatory region, whose mutation reduces in half the expression of a gene of interest can be estimated in 10 (based on data for *lacZ*). Thus, μ_b_ = 10·μ, which means 1 (= μ_b_·〈*N*〉) mutant of this type in the population on average. This frequency may even be higher if we not only consider the mutations in the *lac* promoter, but also the mutations in the coding region, or affecting the activity of its regulators (*e.g*., CRP; Kinney *et al*., 2010).

In addition, for the *lacZ* gene, its duplication creation rate is of μ_c_ = 3·10^−4^ dup./gene/gen., and its duplication deletion rate of μ_d_ = 4.1·10^−4^ - 4.4·10^−2^ -/gene/gen. (Reams *et al*., 2010; Reams *et al*., 2012). In absence of lactose, duplications are neutral (*S* = 0), which means, on average, a duplication frequency in the population of 0.68% - 42% [= μ_c_ / (μ_c_ + μ_d_)]. By contrast, in presence of lactose, duplications are deleterious (*S* ≈ -28%), and then the average duplication frequency is of 0.09% - 0.11 % [= μ_c_ / (μ_c_ + μ_d_ - *S*)]. Note that the deletion rates are difficult to estimate experimentally, as this requires starting from a genotype with new-born (mostly unstable) duplications, albeit they are essential to properly understand the fixation process.

## ACKNOWLEDGEMENTS

This work was supported by grants BFU2015-66894-P (to G.R.) and BFU2015-66073-P (to M.A.F.) from the Spanish Ministry of Economy (MINECO/FEDER).

## CONTRIBUTIONS

GR conceived the study. GR performed the research. GR and MAF analyzed the results. GR wrote the paper with input of MAF.

